# Array-based analysis of SARS-CoV-2, other coronaviruses, and influenza antibodies in convalescent COVID-19 patients

**DOI:** 10.1101/2020.06.15.153064

**Authors:** Daniel J. Steiner, John S. Cognetti, Ethan P. Luta, Alanna M. Klose, Joseph Bucukovski, Michael R. Bryan, Jon J. Schmuke, Phuong Nguyen-Contant, Mark Y. Sangster, David J. Topham, Benjamin L. Miller

## Abstract

Detection of antibodies to upper respiratory pathogens is critical to surveillance, assessment of the immune status of individuals, vaccine development, and basic biology. The urgent need for antibody detection tools has proven particularly acute in the COVID-19 era. We report a multiplex label-free antigen microarray on the Arrayed Imaging Reflectometry (AIR) platform for detection of antibodies to SARS-CoV-2, SARS-CoV-1, MERS, three circulating coronavirus strains (HKU1, 229E, OC43) and three strains of influenza. We find that the array is readily able to distinguish uninfected from convalescent COVID-19 subjects, and provides quantitative information about total Ig, as well as IgG- and IgM-specific responses.

## Introduction

The ongoing SARS-CoV-2 pandemic has had enormous costs in terms of lives lost, impacts to quality of life, and the global economy. In order to reduce the impact of this virus over time, it is widely recognized that assessing human immunity to SARS-CoV-2 will have a critical role to play in safeguarding public health. Detection of anti-SARS-CoV-2 antibodies can provide a clearer understanding of the actual infection rate of an area. It is also hypothesized that those with antibodies to SARS-CoV-2 are protected from reinfection, and therefore able to return to work without health concern.^1^ Indeed, emerging data indicates that COVID-19 patients can achieve a robust immune response.^2,3^ Although recent anecdotal reports suggest that reinfection can occur, even here an ability to measure and quantify coronavirus antibodies will be important to assess whether reinfection probability correlates with low antibody titer, or potentially with antibody responses to some antigens and not others, or correlates with the presence or lack of immunity to other respiratory pathogens. Analytical tools for monitoring antibody responses to specific antigens are also of obvious importance in the development of new vaccines and for understanding fundamental aspects of disease course.^4^

In response to this need, industry and academia have both risen to the challenge,^5^ beginning with several reports on SARS-CoV-2 ELISA assays in late February and early March 2020.^6,7,8,9^ However, most tests that have been reported to date rely on the response of single antigens. Concerns have also been raised regarding the accuracy of some tests.^10,11^ There is a need to understand the preponderance of SARS-CoV-2 antibodies in the general population. It is also critical to study the human immune response following infection. Singleplex tests, whether rapid or implemented in a clinical laboratory, are beginning to provide this information. What they do not provide, however, is a broader understanding of the human immune response to SARS-CoV-2 infection, or illuminate potential relationships between COVID-19 infection and previous infections (and immunity to) other respiratory viruses including circulating coronaviruses that cause the common cold. To address these goals, multiplex analytical techniques are required. A bead-based multiplex immunoassay for six coronaviruses infecting humans (pre-SARS-CoV-2) has been reported,^12^ and more recently a 4-plex assay on the Quanterix platform focused on SARS-CoV-2 antigens has been described.^13^ Despite these advances, there remains a significant need for analytical methods able to rapidly quantify antibodies not only to SARS-CoV-2, but also to other coronaviruses, and other pathogenic viruses. Most importantly, these must be able to discriminate among responses to different closely related viruses and different antigens from the same virus. To address this need, we have developed a prototype 15-plex array on the Arrayed Imaging Reflectometry (AIR) platform.

AIR is a label-free multiplex sensor method in which the surface chemistry and deposition of capture molecules to form a microarray on a silicon chip are carefully controlled such that s-polarized HeNe laser light at a 70.6° incident angle to the chip undergoes total destructive interference within the surface film.^14^ Binding to any probe spot on the array degrades the antireflective condition in proportion to the amount of material bound, yielding an increase in the reflected light as observed by a CCD camera. By comparing the intensity of the reflected light to an experimentally validated model, the thickness change for each spot, and therefore the quantity of each analyte in the sample, may be precisely and sensitively determined.^15^

We have previously reported the utility of influenza antigen arrays fabricated on the AIR platform for assessment of anti-influenza antibodies in human, animal, and avian serum,^16,17^ both as a tool for viral surveillance and for assessment of the efficacy of a candidate vaccine. We have also demonstrated that AIR is scalable at least to 115-plex assays, used for discriminating different influenza virus serotypes.^18,19^ We therefore anticipated that the platform would be useful as a way to quantify anti-SARS-CoV-2 antibodies, antibodies to other coronaviruses including circulating (“common cold”) strains, and other respiratory pathogens including influenza. Here, we discuss the development and testing of a mixed coronavirus / influenza antigen panel on AIR, and its application to analyzing the coronavirus antibody profile of a cohort of convalescent COVID-19 patients and subjects of unknown disease status.

## Methods

### Material sources

For AIR assays, SARS-CoV-2, SARS-CoV, MERS, and Influenza Type A and B antigens were obtained from Sino Biological, Inc., and are described in more detail below. Most antigens were supplied as lyophilized material and reconstituted at the recommended concentrations using 18-MΩ water, while the remaining antigens were supplied frozen on dry ice. PBS-ET was prepared as phosphate buffer (10 mM monobasic sodium phosphate, 10 mM dibasic sodium phosphate, 150 mM NaCl) with 0.02% w/v Tween-20 and 5 mM EDTA. Amine-reactive substrates for fabrication of AIR arrays were provided by Adarza BioSystems, Inc. For ELISA assays, SARS-CoV-2 full-length spike and RBD were produced in-house using a mammalian expression system,^20,21^ as was influenza A/H1N1/California 2009 hemagglutinin. SARS-CoV2 nucleocapsid expressed in mammalian cells was obtained from RayBiotech, while HCoV-229E and HCoV-OC43 spike proteins (baculovirus-expressed) were obtained from Sino Biological. Tetanus toxoid (TTd) was obtained from Calbiochem.

### Antigen probe formulation

Prior to microarray fabrication, antigens were buffer-exchanged and concentrated using Amicon centrifugation filters (EMD Millipore) into phosphate buffer at pH 5.8 and pH 7.2 prior to use. During development, several printing concentrations and/or solution pH values of each antigen were tested, along with sugar additives (glycerol, trehalose) in order to optimize spot uniformity and morphology as well as initial probe thickness.^22^ Antigen concentrations and pH values printed in the final arrays to generate all data in this publication are shown in Table 1.

**Table 1.** Formulation parameters for printed antigen solutions.

| Array ID | Antigen | Probe Concentration ($\mu\text{g/mL}$) | pH | Additives |
| --- | --- | --- | --- | --- |
| F1 | α-FITC 650 | 650 | 7.2 | 2.5% Trehalose |
| F2 | α-FITC 400 | 400 | 7.2 | 5% Glycerol |
| F3 | α-FITC 400 + 5% glycerol | 400 | 5.8 | 2.5% Trehalose |
| 1 | Human coronavirus spike glycoprotein, HKU isolate (aa 1-760, His tag) | 200 | 5.8 | 2.5% Trehalose |
| 2 | MERS-CoV (nCoV/Novel coronavirus) Spike Protein S1 (aa 1-725, His Tag) | 270 | 7.2 | 2.5% Trehalose |
| 3 | MERS-CoV (nCoV/Novel coronavirus) Spike Protein fragment (RBD, aa 367-606, His Tag) | 300 | 7.2 | 2.5% Trehalose |
| 4 | SARS-CoV Spike/S1 Protein (S1 Subunit, His Tag) | 205 | 5.8 | 2.5% Trehalose |
| 5 | Human SARS Coronavirus Spike Protein (Receptor Binding Domain, His tag) | 141 | 7.2 | 2.5% Trehalose |
| 6 | NCP-CoV (2019-nCoV) Spike Protein (S1+S2 ECD, His tag) | 400 | 7.2 | 2.5% Trehalose |
| 7 | SARS-CoV-2 (2019-nCoV) Spike Protein (S2 ECD, His tag) | 150 | 5.8 | 2.5% Trehalose |
| 8 | SARS-CoV-2 (2019-nCoV) Spike Protein (S1 Subunit, His tag) | 300 | 7.2 | 2.5% Trehalose |
| 9 | 2019-nCoV Spike Protein (RBD, His Tag) | 400 | 7.2 | 2.5% Trehalose |
| 10 | Human coronavirus (HCoV-229E) Spike Protein (S1+S2 ECD, His Tag) | 200 | 7.2 | 2.5% Trehalose |
| 11 | Human coronavirus (HCoV-OC43) Spike Protein (S1+S2 ECD, His Tag) | 200 | 7.2 | 2.5% Trehalose |
| 12 | Influenza B/Brisbane/2008 HA | 100 | 5.8 | 2.5% Trehalose |
| 13 | Influenza A/California/2009 (H1N1) HA | 100 | 5.8 | 2.5% Trehalose |
| 14 | Influenza A/Wisconsin/2005 (H3N2) HA | 100 | 5.8 | 2.5% Trehalose |

### Preparation of arrays

Arrays were printed on amine-reactive silicon oxide substrates (Adarza BioSystems, Inc.) using a Scienion SX piezoelectric microarrayer (Scienion, A.G.) with spot volumes of approximately 300 pL. Six spots were printed for each antigen, the final layout of which is shown in Figure 1. The number of spots arrayed was not critical to robust analytical performance or statistical analysis. Each spot consists of approximately 300 pixels when imaged by the CCD in an AIR chip reader (Adarza BioSystems, Inc.), with each pixel representing a discrete interrogation of a unique probe surface region. Therefore, averaging these pixel values together produces an inherently reliable measure of analyte-to-probe response. Dilutions of polyclonal anti-fluorescein (anti-FITC, Rockland Inc.), were printed as negative intra-array controls. After printing, chips were mounted onto adhesive strips at appropriate spacing for 96-well plates, and then placed into 50 mM sodium acetate buffer (pH 5) for 5 minutes. Next, a 1.5% BSA solution was added to each well resulting in a final BSA concentration of 0.5% to passivate the remaining amine-reactive surface functionality. After blocking for 20 minutes, the chips were transferred to new wells containing 20% fetal bovine serum (Gibco) in PBS-ET as a secondary block, and incubated for 40 min. This step was required to reduce nonspecific binding from human serum at the assay endpoint. The chips were then rinsed briefly (5 min) in new wells containing PBS-ET, then transferred to wells containing Microarray Stabilizer Solution (Surmodics IVD). After a 30-minute incubation, the chips were dried at 40 °C in an oven for 60 min. This last step renders the sensors shelf-stable, until use in assays performed later.

**Figure 1:**
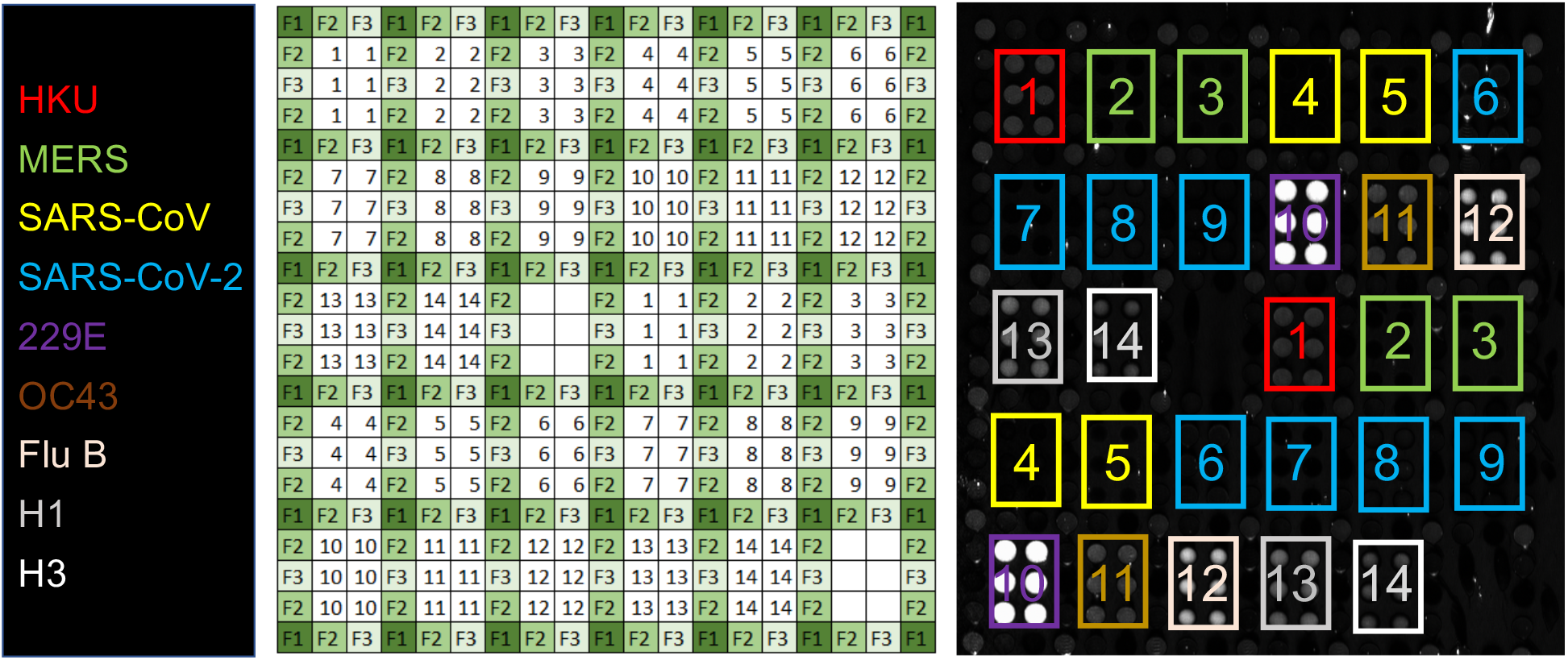
AIR assay for antibodies to respiratory viruses. For each antigen, six replicate spots are printed in two different locations on the chip. Each group of six spots is surrounded by negative control reference spots (anti-FITC). Blank (background) areas are included as additional negative controls. Key: **1:** human coronavirus (HKU isolate) spike glycoprotein, aa 1-760; **2:** MERS-CoV spike glycoprotein, S1 domain; **3:** MERS-CoV spike glycoprotein, receptor binding domain (RBD); **4:** SARS-CoV spike glycoprotein, S1 domain; **5:** SARS-CoV spike glycoprotein, RBD; **6:** SARS-CoV-2 spike glycoprotein, S1+S2 ECD; **7:** SARS-CoV-2 spike glycoprotein, S2 ECD; **8:** SARS-CoV-2 spike glycoprotein, S1 domain; **9:** SARS-CoV-2 spike glycoprotein, RBD; **10:** human coronavirus (HCoV-229E isolate) spike glycoprotein, S1+S2 ECD; **11:** human coronavirus (HCoV-OC43 isolate) spike glycoprotein, S1+S2 ECD; **12:** influenza B/Brisbane/2008 hemagglutinin; **13:** influenza A/California/2009 (H1N1) hemagglutinin; **14:** influenza A/Wisconsin/2005 (H3N2) influenza. **F1**, **F2**, and **F3** are derived from spotting three different dilutions of anti-FITC. The image at right is a representative array exposed to Pooled Normal Human Serum (PNHS) at a 1:4 dilution.

### AIR Assay protocol

A sample diluent consisting of a proprietary buffer (Adarza BioSystems, Inc.), 20% fetal bovine serum (FBS), and in some cases (polyclonal antibody titrations; discussed later), pooled normal human serum (PNHS; 10% v/v, Innovative Research) was used to dilute monoclonal and polyclonal antibodies as well as donor human serum samples to appropriate standard concentrations. Wells in a 96-well plate to be used for target solutions were first treated with a preblock solution for 40 minutes (10 mg/mL BSA in 1x mPBS-ET, pH 7.4, 0.2 μm sterile filtered). This was then pipetted out, and replaced with target solution. Arrays were incubated with target solutions overnight at 4 °C with orbital agitation on a microtiter plate shaker (500 RPM). Chips were then removed after target analyte exposure and rinsed by transferring to wells containing PBS-ET for 5 min, twice. After washing, chips were rinsed under flowing 18-MΩ water and dried under a stream of nitrogen. Finally, the substrates were imaged using a prototype AIR Reader and internally developed imaging software at several integration times with dark field subtraction. AIR assays run using the commercial Adarza Ziva instrument were incubated 30 minutes at room temperature before undergoing automated processing and imaging by the instrument.

### AIR antibody class assessment

To determine whether samples consisted of a primarily IgG- or IgM-based response, we performed an experiment including a secondary antibody incubation step. After overnight primary incubation with a selection of both positive and negative samples (based on prior AIR assays), chips were removed from sample wells and washed twice with PBS-ET. Then chips for each sample were placed in wells containing either anti-human IgG, IgM, or 20% FBS as a negative control. Each of these conditions was produced in duplicate. Secondary antibodies were diluted to 1 μg/mL for both goat α-hIgG (Jackson Immunoresearch) and rabbit α-hIgM (Rockland, Inc.) in Adarza diluent. After one hour of incubation with secondary antibodies at room temperature, chips were washed twice for 5 minutes in PBS-ET, then rinsed with water and dried with nitrogen as before.

### Human samples

Whole blood was drawn via venipuncture, allowed to clot at ambient temperature for 30 minutes, and then centrifuged at 1200 x g for 15 minutes. Serum was drawn off via pipette, aliquoted, and stored at −80 °C prior to use. Sera were drawn under a protocol approved by the University of Rochester Medical Center Institutional Review Board. Data acquired by Adarza BioSystems, Inc., used samples obtained from commercial sources (pre-COVID-19 negative samples: Innovative Research, Inc.; COVID-19 convalescent samples: Discovery Life Sciences).

### Data analysis

AIR images were analyzed using the Adarza ZIVA data analysis tool. Probe spots with major defects or debris were manually flagged and eliminated, and minor defects in spot quality were automatically identified and excluded from the median intensity measurement. The median intensity values were converted to median thickness values using a best-fit line to an experimentally derived reflectance model.^14^ Then, the median thickness values were further processed in Microsoft Excel as described below, and are referred to simply as “thickness” hereafter.

While anti-FITC spots were designed to serve as an intra-chip normalizer, these were not used as such due to the unexpected presence of anti-goat IgG antibodies in some single donor human serum samples. Therefore, the blank area served an intra-chip normalizer to mitigate any variation in the reactivity of the surface chemistry between AIR chips. The thickness of the blank area was subtracted from the thickness of each probe spot to produce “normalized thickness” values for each probe spot.

All of the normalized thickness values across replicate chips (n=2) were averaged together (maximum of n=24 probe spots) for each antigen, and the standard deviation was calculated. The average thickness for each antigen in the fetal bovine serum (FBS) control was subtracted from the average thickness obtained for each antigen in each subject sample to produce the “normalized thickness change (Δ Thickness).” In the case of the polyclonal antibody titration, the control chip was incubated in a matrix of FBS and PNHS.

### ELISA assay

Serum IgG titers specific for SARS-CoV-2 proteins and selected non-coronavirus proteins were determined by ELISA as described previously.^23^ In brief, NUNC MaxiSorp 96-well ELISA plates (Invitrogen, Carlsbad, CA) were coated with optimized concentrations of coating reagents at least one day prior to the assay. After blocking plates with 3% BSA/PBS for 1 h, serial 3-fold dilutions of serum samples in ELISA diluent (0.5% BSA/0.05% Tween-20/PBS) were added and incubated for 2 h. Antigen-specific IgG was detected by addition of alkaline phosphatase-conjugated anti-human IgG (clone MT78; Mabtech, Cincinnati, OH), followed by p-nitrophenyl phosphate substrate. Well absorbance was read at 405 nm after color development. Human serum standards were used to assign weight-based concentrations of antigen-specific IgG as previously described, with the limit of assay sensitivity set at 0.5 μg/mL for all antigens.^23,24^

## Results

### Initial array qualification

Arrays were initially qualified using commercial mono- and polyclonal antibodies (Sino Biological) doped in PNHS. PNHS alone produced strong signals to circulating (common cold) coronaviruses HKU, OC32, and 229E, as well as to the three influenza hemagglutinins on the array [B/Brisbane/2008, A/California/2009 (H1N1), A/Wisconsin/2005 (H3N2)]. This is as expected given the prevalence of these viruses in the general population. Addition of an anti-SARS-CoV-2 polyclonal antibody raised against the SARS-CoV-2 spike protein receptor binding domain (RBD) at 1 μg/mL produced a strong signal on all three RBD-containing antigens (S1 + S2 ECD, S1, and RBD). Overall response to the polyclonal antibody was well-behaved, and titrated to zero as expected (Figure 2C and D). Quantitative data are presented in Ångstroms of build. At the highest concentrations, significant cross-reactive binding to the HCoV-229E spike protein was observed, as well as some binding to the HCoV-OC43 spike protein and MERS S1. Calculated limits of detection^25^ for these data were 43.3 ng/mL (SARS-CoV-2 S1 + S2 ECD), 40.7 ng/mL (SARS-CoV-2 S1), and 25.1 ng/mL (SARS-CoV-2 RBD). However, these should be viewed as provisional, and subject to optimization.

**Figure 2:**
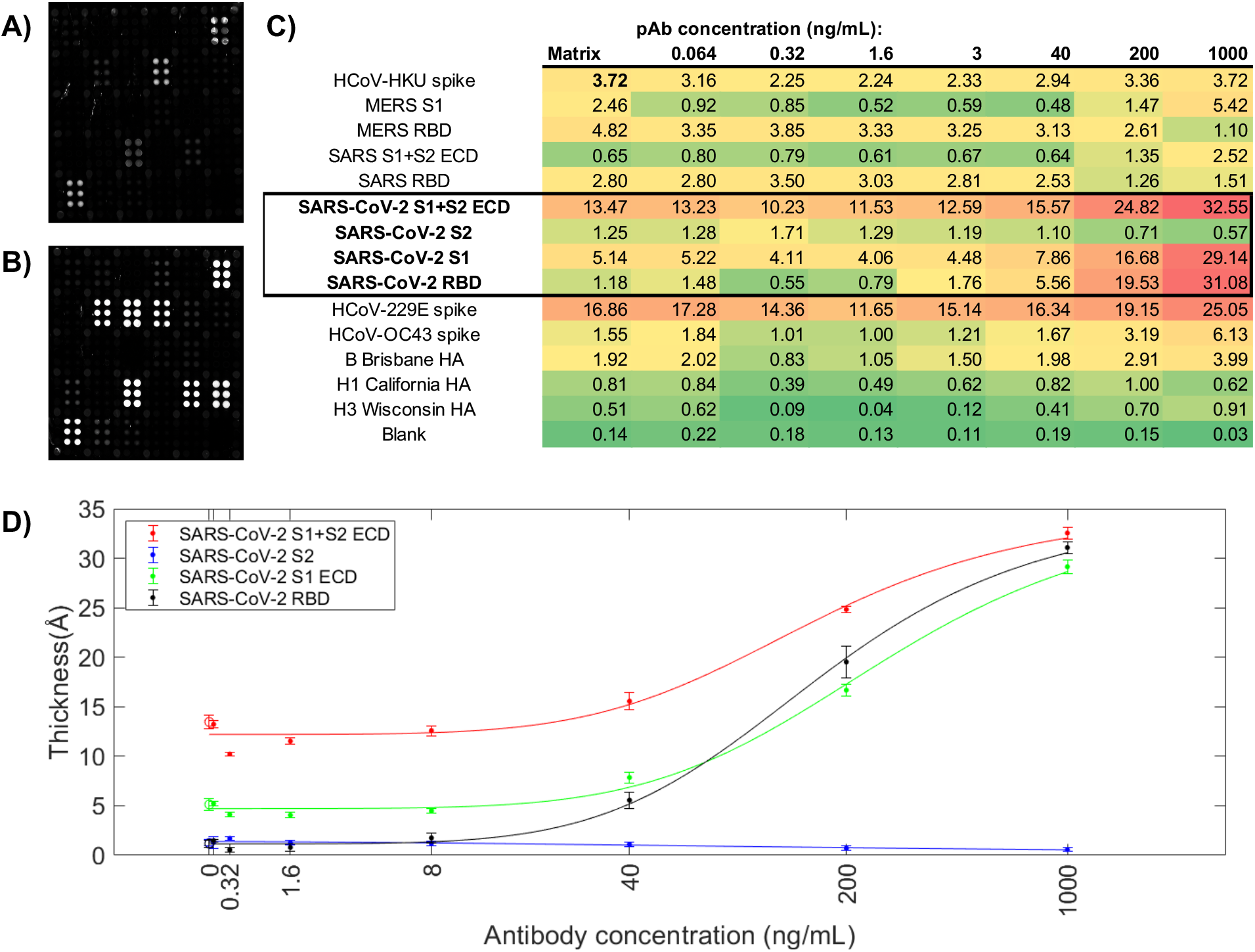
Response of a commercial anti-SARS-CoV-2 rabbit polyclonal antibody (pAb) on the array. (A) array exposed to array exposed to 20% FBS + 10% PNHS; (B) array exposed to 1 μg/mL anti-SARS-CoV-2 pAb in 20% FBS + 10% PNHS. Strong responses to SARS-CoV-2 S1+S2 ECD, S1, and RBD are observed, as well as smaller cross-reactive responses to HCoV-229E, HCoV-OC43, and MERS spike proteins; (C) quantitative data for the titration. Concentrations of pAb are provided at the top of each column in ng/mL; response values at each concentration for each antigen are provided in Ångstroms of build. (D) Titration curves for the four SARS-CoV-2 antigens with standard deviation of replicate probe spots at each concentration.

With initial qualification of the array completed, we turned our attention to examination of a series of human serum samples. At the outset of our study, few samples from known COVID-19 patients were available. Thus, the first individual donor samples constituted a small group of healthy individuals with no known COVID-19 diagnosis. Later, a set of 15 samples were obtained from convalescent COVID-19 patients at least 14 days out of active disease, and acquired via the University of Rochester Medical Center's Healthy Donor protocol. Figure 3 shows a comparison of array images obtained for FBS, PNHS, and a representative convalescent COVID-19 patient. Strong responses to SARS-CoV-2 antigens are readily visible in the array exposed to the COVID-19 patient's sample. Differences in the response of the known positive sample to non-SARS-CoV-2 antigens relative to PNHS are readily visible, and discussed in more detail in the context of quantitative analysis, below.

**Figure 3:**
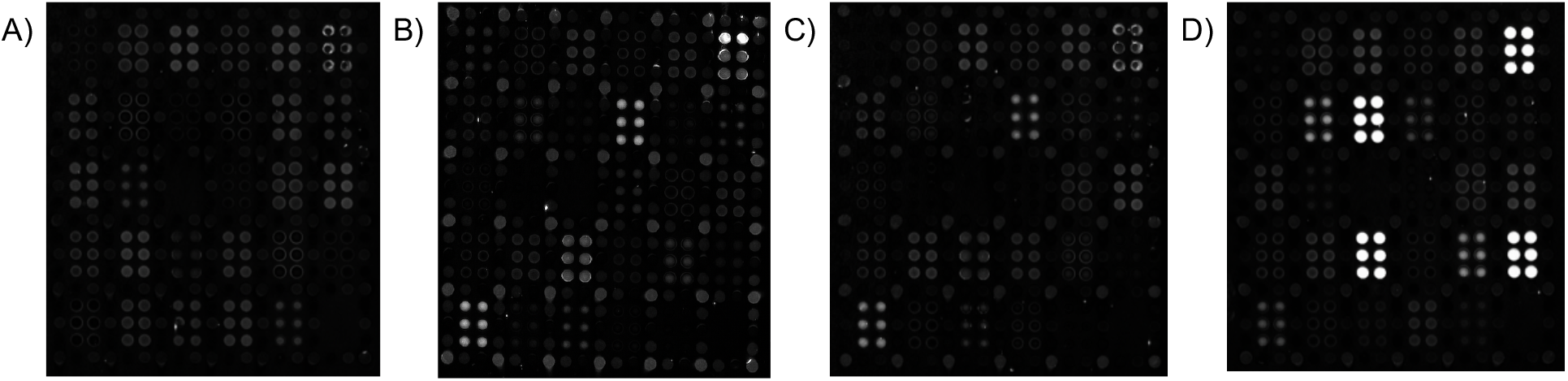
Representative AIR array images (100 ms exposures) of (A) 5% FBS; (B) 10% PNHS; (C) a negative single-donor sample, and (D) one convalescent serum sample. Strong responses to SARS-CoV-2 antigens are readily observed in (D), but not in (A), (B), or (C). In each case, samples were diluted 1:20 in Adarza diluent, and incubated with the arrays overnight at 4 °C. See Figure 1 for key to the array. All arrays in this figure were imaged at an exposure of 100 ms.

Quantification of responses was conducted for all arrays as described in the *methods* section, with results presented for 1:5 dilution samples in Figure 4 (A). Most samples from convalescent COVID-19 patients yielded robust responses to at least one SARS-CoV-2 antigen. Small negative “build” values indicate subtle difference in sample matrix relative to the control, and can be discounted. Sample HD2135 produced minimal signal on all coronavirus antigens. After unblinding the clinical details of these subjects, we discovered that this sample derived from a person self-reporting COVID-19, but with no recorded positive PCR test for SARS-CoV-2. Sample HD2146 also was unreactive with SARS-CoV-2 antigens, despite having experienced COVID-like symptoms and receiving a positive PCR result. Similar results were obtained via ELISA with this sample (*vide infra*), suggesting the discrepancy was not technique dependent. Samples were also run at a 1:20 dilution; as shown in Figure 4 (B), results were similar but not identical to the 1:5 dilution results. It is likely that nonspecific binding by serum proteins influences the results from more concentrated samples, and thus future experiments will focus on 1:20 and higher dilutions.

**Figure 4:**
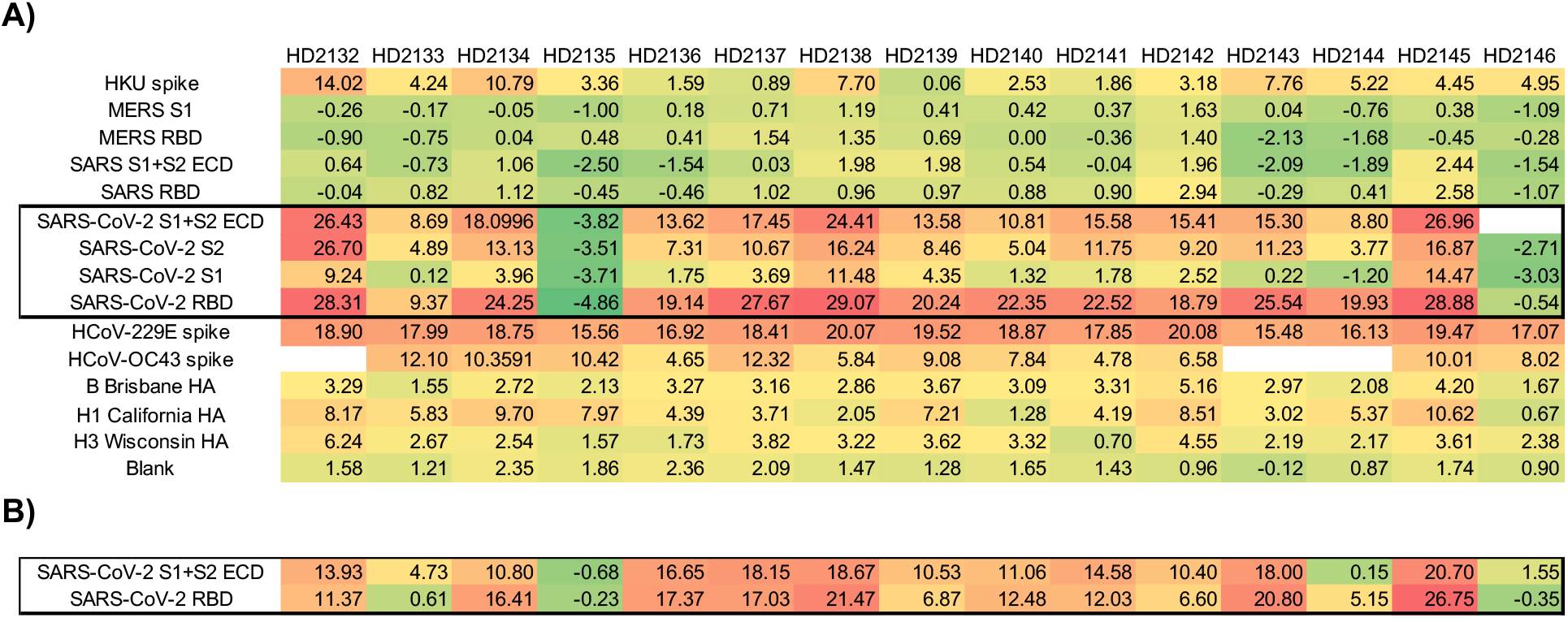
AIR results from convalescent COVID-19-positive subjects. Empty cells indicate unreadable spots. (A) Each sample was diluted 1:5 in Adarza diluent, and incubated with the array overnight at 4 °C. (B) Selected antigen results for samples run at 1:20 dilution, and incubated with the array overnight at 4 °C. All values reported are in Ångstroms of build relative to an FBS control.

Convalescent serum array responses were compared to an ELISA assay (Figure 5). As ELISA values were all IgG-specific, and AIR data discussed thus far (obtained in a “label-free” mode) was a combination of IgG and IgM-specific responses, these results would not be expected to match precisely. Differences in the expression system used for antigen production (baculovirus for commercial antigens used in AIR; HEK 293T cells used for antigens used in the ELISA assays) could also lead to differences. However, overall trends for SARS-CoV-2 antigens correlate well, as shown in Figure 6. To provide further detail with regard to the response, AIR assays were run using secondary anti-IgG and anti-IgM antibodies to determine class-specific responses for a subset of samples (Figure 7).

**Figure 5:**
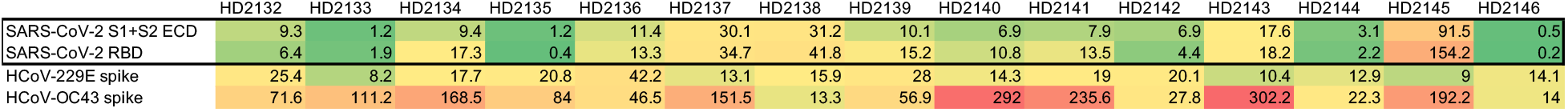
IgG-specific ELISA results for convalescent COVID-19 subjects. All values are in μg/mL.

**Figure 6:**
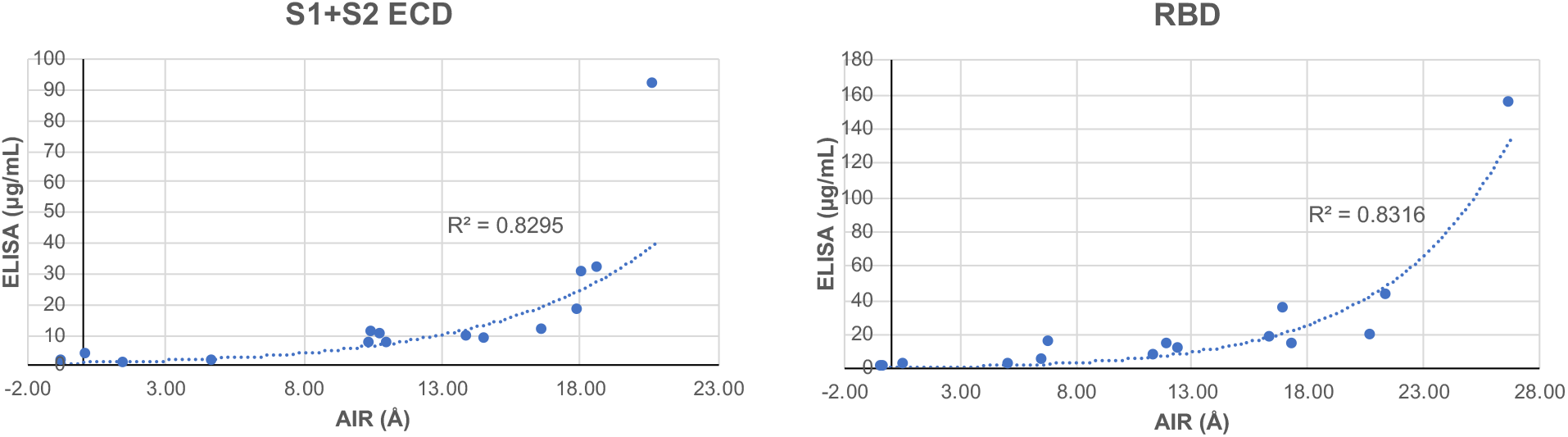
Correlation of AIR and ELISA data for SARS-CoV-2 S1+S2 ECD (left) and RBD (right). Exponential trend lines and associated R^2^ values are indicated.

**Figure 7:**
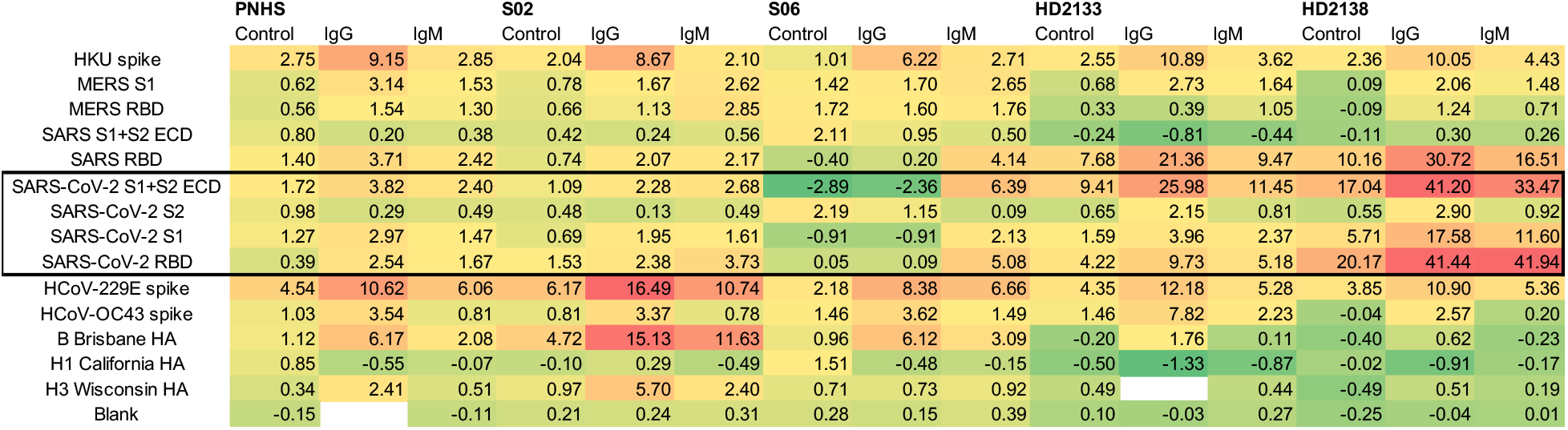
Determination of class-specific responses for a subset of COVID-19-negative (S02, S06) and convalescent COVID-19-positive (HD2133, HD2138) subjects. All values reported are in Ångstroms of build relative to an FBS control.

Finally, a laboratory assay is most valuable to others if it can be scaled up by a commercial facility in a manner enabling its broad distribution. To address that issue, Adarza BioSystems printed analogues of our arrays for use in the ZIVA system, an automated version of AIR employing a consumable with a low volume requirement (25 μL). Fifteen samples from convalescent COVID-19 patients were tested using a 30-minute room-temperature incubation, and compared with 16 single-donor samples acquired prior to the outbreak of COVID-19. As was the case with assays run using the laboratory AIR assay, analysis using the ZIVA system readily discriminated between negative and convalescent samples (Figure 8). Three putative convalescent COVID-19 samples gave responses on all SARS-CoV-2 antigens that were below the threshold for a positive response (two standard deviations above the average of the 16 negative samples). This is analogous to the AIR and ELISA results obtained for sample HD2146, as described above. The remaining 12 convalescent samples gave strong responses on at least one SARS-CoV-2 antigen, with many responding strongly to both RBD and S2 (Figure 8).

**Figure 8:**
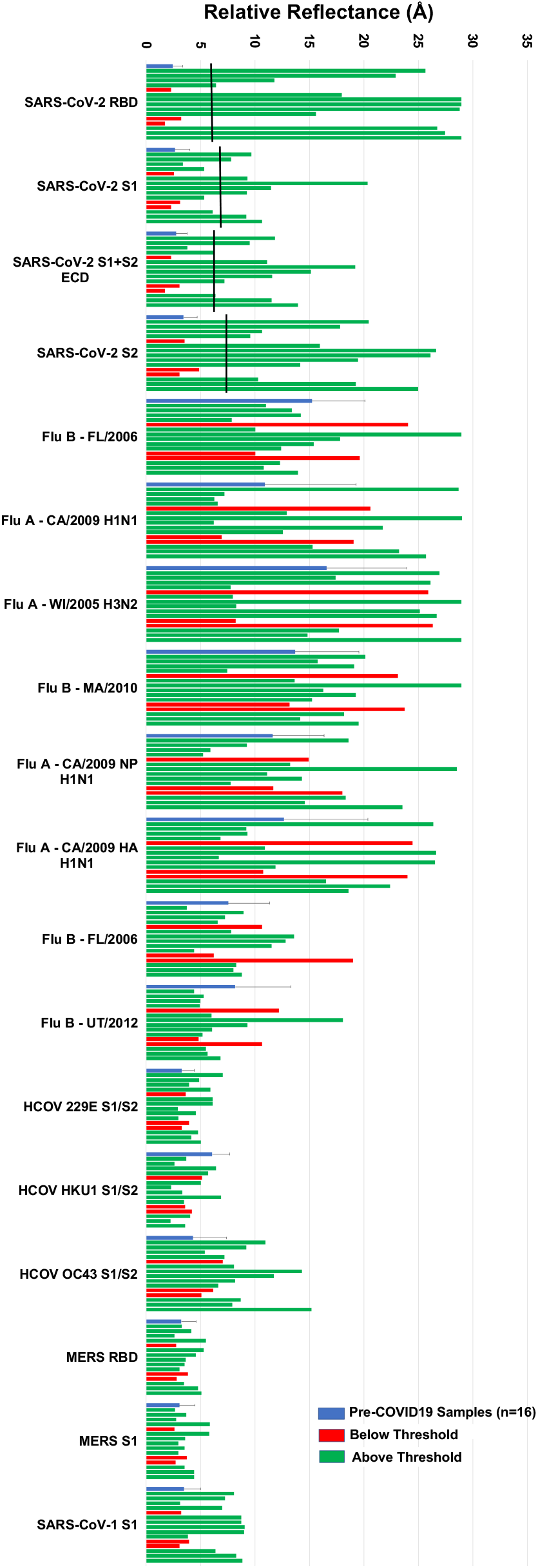
Results from the Adarza Ziva system for pre-COVID-19 serum samples and single-donor samples from convalescent COVID-19 (PCR-positive) subjects. Pre-COVID-19 single-donor results were averaged (blue bars). Black bars indicate threshold positive values, calculated as two standard deviations above the average negative (pre-COVID-19) signal. Red bars indicate PCR+ individuals yielding signals below the threshold on all SARS-CoV-2 antigens, while green bars indicate signals from single-donor convalescent COVID-19 samples with at least one SARS-CoV-2 antigen response above threshold.

## Discussion

Health and disease result from many factors, including the overall landscape of a person's immune system. As such, methods for profiling antigen-specific antibody titers to a range of diseases in addition to the disease of primary current interest are of utility when studying the disease. To that end, we have presented preliminary data on a 15-plex array on the AIR platform, developed in response to the need to study SARS-CoV-2 but incorporating antigens for other coronaviruses and influenza. Responses to SARS-CoV-2 antigens on the array effectively discriminated between serum samples from uninfected and COVID-19 convalescent subjects, with generally good correlation to ELISA data. Follow-up assays demonstrated that exposure of the arrays to anti-IgG and anti-IgM antibodies enabled discrimination of antibody isotype.

An important aspect of this work is the ability to evaluate anti-SARS-CoV-2 immunity in the context of the individual's overall immune landscape. Because available chip real estate allows for substantial expansion of the multiplex capability of the array, in ongoing efforts we will add additional antigens for other strains of influenza (by analogy to our previous work^17^), as well as other upper respiratory infections such as respiratory syncytial virus and metapneumovirus. Other coronavirus antigens including nucleocapsid (N) are also likely candidates for addition to the array, as they are known to produce an immune response (as seen in the ELISA results, for example). Thus, the flexibility of the AIR platform will prove useful not only in the current pandemic, but as other viruses inevitably emerge.

## Acknowledgement

We thank Alicia Papalia for assistance with human samples, and Florian Krammer for the generous donation of plasmids for SARS-CoV-2 antigen production (S1 + S2 ECD and RBD). Support from the University of Rochester Department of Dermatology and Adarza BioSystems are gratefully acknowledged. This project has been funded in whole or in part with Federal funds from the National Institute of Allergy and Infectious Diseases, National Institutes of Health, Department of Health and Human Services, under CEIRS Contract No. HHSN272201400005C.

